# Low-cost gold-leaf electrode as a platform for *Escherichia coli* immuno-detection

**DOI:** 10.1101/2022.12.14.520406

**Authors:** Ivana Podunavac, Manil Kukkar, Vincent Léguillier, Francesco Rizzotto, Zoran Pavlovic, Ljiljana Janjušević, Vlad Costache, Vasa Radonic, Jasmina Vidic

## Abstract

Gold electrodes are one of most prevalent substrates in electrochemical biosensors because they can be easily and highly efficiently functionalized with thiolated biomolecules. However, conventional methods to fabricate gold electrodes are costly, time consuming and require onerous equipment. Here, an affordable method for rapid fabrication of an electrochemical immune-sensor for *Escherichia coli* detection is presented. The gold electrode was generated using 24-karat gold leaves and low-cost polyvinyl chloride adhesive sheets covered with an insulating PTFE layer. The gold-leaf electrode (GLE) was patterned using laser ablation and characterized by cyclic voltammetry, electrochemical impedance spectroscopy, scanning electronic microscopy, contact angle and 3D profiling. The GLEs were modified by a self-assembled mercaptopropionic monolayer, followed by surface activation to allow binding of the specific anti-*E. coli* antibody via carbodiimide linking. The biosensor showed a detection limit of 2 CFU/ml and a linear dynamic range of 10 – 10^7^ CFU/ml for *E. coli* cells. No false positive signals were obtained from control bacteria. The obtained results demonstrated suitability of GLE for use in biosensors with high reliability and reproducibility. It is foreseeable that our work will inspire design of point-of-need biosensors broadly applicable in low-resource settings.

## 1. Introduction

Gold electrodes are frequently used in biosensors due to excellent electrical conductivity, outstanding mechanical stability, and easy functionalization of gold through the formation of strength gold–thiol bonds (Bobrinetskiy et al. 2021; Sui and Zorman 2020; Vidic et al. 2007). Electrochemical biosensors have been developed to detect and monitor an array of biomarkers (DNA, RNA, antigens, antibodies) for applications ranging from water and food quality to human health conditions. However, although electrochemical biosensors are easy to use and can be connected to low-cost and hand-held instrumentation only a limited number has been commercialized. In part, this comes from the challenges related to gold electrode fabrication.

Traditionally, gold electrodes fabricated by photolithography, or shadow mask lithography suffer from disadvantages such as laborious and complex assembly processes, use of costly and harmful chemicals and necessity for specialized laboratories and cleanrooms (Sui and Zorman 2020), which are inaccessible to many. Other methods, such as electroplating and vacuum deposition have a high cost. In addition, they need to be associated with an etching step, where deposited materials are selectively removed and converted into waste that in some cases has to be treated. Inkjet and screen printing depositions have risen in importance as alternative affordable fabrication methods to traditional methods for electrode fabrication. Inkjet-printed electrodes are generated by applying a gold nanoparticle ink to a substrate or previously deposited layers (Singh et al. 2010). Effective structures with fine patterns can be obtained with a small amount of nanoparticles, without the need for shadow masks and using simple equipment. However, the inevitable tendency of dissolved nanoparticles to form coffee-ring-like structures due to surface-tension-driven transport and the flow properties of interacting droplets with the surface may produce a non-uniform film, which decreases device performance. To avoid this, gold ink can be doped with other materials and applied to a substrate through a patterned mask such as the case of commercialized disposable gold screen-printed electrodes (SPEs) (Bernalte et al. 2013). The heterogeneity and irreproducibility reported with SPEs probably originated from the doping material that can reduce the efficiency of electrode functionalization or interfere with sensing.

Very low-cost gold leaf sheets of nanometer thickness made of almost pure gold are not a frequently used material for making electrochemical sensors because of the difficulty in sheets handling and their fragility (Prasertying et al. 2021). Up to today, only a few reports present pure gold leaf as electrochemical sensors: Matsui et al., developed a microband gold-leaf electrode for glucose sensing (Matsui et al. 2017), Zamani et al., produced a gold-leaf electrode for sensing DNAse I activity (Zamani et al. 2021), and Prasertying et al., fabricated a planar electrochemical sensor comprising gold-leaf as the working electrode for Pb^2+^-ion detection (Prasertying et al. 2021). In these cases, gold-leaf electrodes were obtained by mounting the leaf between two layers of insulating polyimide, polyester or polyvinyl chloride (PVC) adhesive tape. The upper tape layer had a hole to enable exposure of the electrode surface to the sample solution.

Regarding to healthcare and agri-food sectors, *Escherichia coli* is not only a member of the most relevant foodborne and waterborne pathogens but also a major reservoir of antimicrobial resistance genes. *E. coli* strains are predominantly harmless commensals in the gastrointestinal tract of mammals and birds and can reside, independent of a host, in water, soil and sediments. When adapted to extra-intestinal niches in humans, *E. coli* can cause diseases with a significant burden on health systems globally, such as urinary tract infections, pyelonephritis, sepsis and meningitis (Manges et al. 2019). In addition, *E. coli* is a fecal indicator bacterium in environmental water quality testing (Kotsiri et al. 2019) and an important indicator in food safety assessment (Marin et al. 2021). Various biosensors based on antibodies have been reported for *E. coli* detection in order to provide rapid diagnostics and to improve measures for health protection (Bonnet et al. 2018; Dastider et al. 2013; El Ichi et al. 2014; Tang et al. 2016; Vidic and Manzano 2021; Zhang et al. 2020).

In this paper, we present development of in-house fabricated disposable gold-leaf electrodes (GLEs) using 24-karat gold leaves and demonstrate their analytical performance in the detection of *E. coli* cells. The working electrode was obtained by applying a gold leaf sheet directly on a substrate and patterned using laser ablation without the need for cleanroom fabrication. The biosensor performance was evaluated by cyclic voltammetry (CV) and electrochemical impedance spectroscopy (EIS). The results indicated that gold-leaf electrodes were suitable for biosensing and their application was demonstrated through sensitive immune-detection.

## 2. Materials and Methods

### 2.1. Chemicals

All chemicals were of analytical reagent grade. Solutions were prepared in deionized MilliQ water. Sulfonic acid, mercaptopropionic acid (MPA), bovine serum albumin (BSA), N-hydroxysuccinimide (NHS), N-(3-dimethylaminopropyle)-N-ethyle-carbodiimidehydro-chloride (EDC), potassium ferricyanide (K_3_[Fe(CN)_6_]), potassium ferrocyanide (K_4_[Fe(CN)_6_]), sodium chloride (NaCl), potassium chloride (KCl) and tryptone soya broth (TBS) were all purchased from Sigma-Aldrich (USA) and used as received. Phosphate buffer saline (PBS) containing 140 mM NaCl, 2.7 mM KCl, 0.1 mM Na_2_HPO_4_ and 1.8 mM KH_2_PO_4_, pH 7.4 was used as an electrolyte. A stock solution of PBS (10×) was purchased from Fisher Bioreagents (United Kingdom). Dry lubricant polytetrafluoroethylene (PTFE) spray was purchased from Wurth (Serbia) and PVC foils “ImageLast A4 125 Micron Laminating Pouch” were purchased from Fellowes Brands (Polska). Gold leaves were purchased from Pozlata Dimitrijevic (Belgrade, Serbia). Rabbit polyclonal anti-*E. coli* antibody (ab137967) was purchased from Abcam (USA). SPE DropSens 220AT were purchased from Metrohm (France). The external reference electrode was a CHI 111 Ag/AgCl, 3 M KCl (CH Instruments, USA).

### 2.2. Fabrication and surface characterization of the gold-leaf electrode

A schematic diagram depicting fabrication of a gold-leaf electrode (GLE) is shown in Fig. 1a. To fabricate the electrode, gold leaf foil (80 mm × 80 mm) was placed onto four-layer PVC sticker foil (4×125 μm), after which it was transferred to the foil through contact of the gold leaf with the tape, followed by light pressing at 180°C using Laminator PDA3 330C (PINGDA, China). In order to make the surface of the PVC sticker hydrophobic, PTFE spray was applied before gold deposition. Custom-made gold leaf electrodes were designed and prepared by laser ablation of gold using a Nd:YAG laser Power Line D-100 (Rofin-Sinar, Germany) in hatch mode and at 25.2 A current, 65 kHz frequency and 500 mm/s speed. A 3D optical profilometer and a Huvitz microscope with Panasis bioimaging software (Republic of Korea) was used for both 2D and 3D imaging of the GLE surface after fabrication to determine the surface roughness.

**Figure 1.**
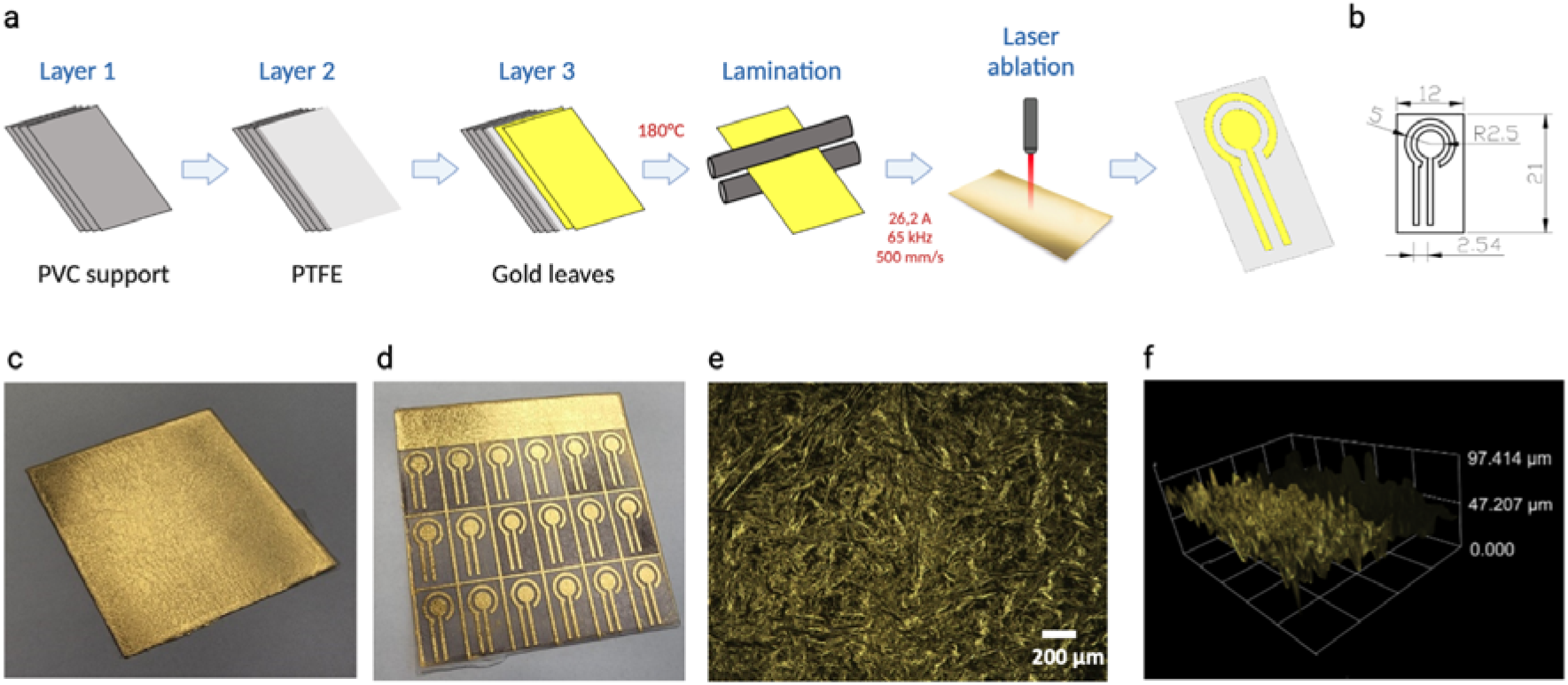
GLE fabrication. (a) Schema of gold-leaf electrode fabrication. (b) Diagram depicting the GLE dimensions. (c) Image of 24-karat gold leaf used for electrode fabrication, dimension 80 × 80 mm. (d) Image of designed electrodes that can be obtained in a single run. (e) 2D profile of the electrode surface. (e) 3D profile of the electrode surface.

Gold-leaf electrodes were cleaned by CV in the presence of 0.5 M H_2_SO_4_ for ten cycles in a potential window of −0.2 to 1.9 V vs. Ag/AgCl at a scan rate of 0.5 V/s. The electrodes were then rinsed with deionized water and dried. To determine the electroactive area of the GLE, cyclic voltammograms were obtained using a 10 mM ferri/ferrocyanide redox couple in PBS. The electroactive surface was calculated according to the Randles-Sevcik equation for reversible electrode process:

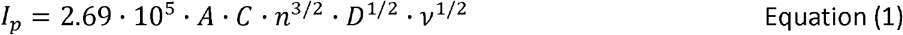

where A is the electroactive area in cm^2^), l_p_ is anodic current peak in A, D is the diffusion coefficient (6.1 · 10^−6^cm^2^/s in case of [Fe(CN)_6_]^4−^ in solution), n is the number of electrons transferred in half-reaction (1 for [Fe(CN)_6_]^4−^), *v* is scan rate (0.5 V/s was used) and C is concentration of [Fe(CN)_6_]^4−^) in mol/cm^2^. Figure S1 shows photo images of GLE and GLE-based devices. The roughness factor (R_f_) of the electrode was estimated by dividing the effective electrode area by its geometric surface area.

### 2.3. Biofunctionalization of the GLE with anti-E. coli antibody

The cleaned GLEs were incubated with 1 mM MPA-ethanolic solution (1 μL) overnight at 4 °C in the dark. A small volume of MPA was used in order to functionalize only the working electrode. The GLEs-MPA were rinsed with ethanol and deionized water and then dried. Subsequently, the functionalized working electrode was exposed to an activation reagent solution (10 μL of 50 mM EDC/50 mM NHS in PBS buffer) for 1 h at room temperature and rinsed with deionized water. The working electrode was then functionalized with 10 μL of 100 μg/ml anti-*E. coli* antibody in PBS at room temperature for 1 h. The GLEs-antibody surface was rinsed with deionized water. Afterwards, the remaining active sites of biofunctionalized GLEs were saturated with 50 μg/ml BSA in PBS at room temperature for 30 min, then rinsed with PBS, followed by deionized water without drying.

### 2.4. Bacterial culturing and quantification

The ATCC 25922 *E. coli*, PY79 *Bacillus subtilis* and ATCC 13048 *Enterobacter aerogenes* bacterial strains used in this study were cultured on TSB agar plates overnight. A single colony was then transferred to a liquid TSB. The culture turbidity was adjusted to match a 0.5 McFarland standard and dilutions (10^8^–10^1^CFU/ml) were performed in PBS. Quantitative bacterial counts were performed in duplicate by plating 0.1 ml aliquot of the limited dilutions (10^3^ and 10^2^ CFU/ml) onto TBS agar plates to confirm the bacterial concentrations.

### 2.5. Electrochemical measurements

Electrochemical measurements were performed with a Potentiostat/Galvanostat/Impedance Analyzer-PalmSens 4 (PalmSens BV, Netherlands), connected to a personal computer equipped with PSTrace 5.8 software. A commercial Ag/AgCl electrode was used as the reference electrode in CV measurements (Fig. S1). CV measurements with ferro-ferricyanide redox probe in PBS were performed in the potential range from −0.3 V to 0.8 V, with the scan rate 0.5 V/s. EIS, which measures both the resistive and capacitive properties of materials via the perturbation of a system at equilibrium with a small sinusoidal excitation signal, was modeled with equivalent circuit models enabling interpretation of the electrical properties of the electrical double layer. Impedance measurements were carried out with the ferro-ferricyanide redox probe prepared in PBS at the room temperature over a frequency range from 1 Hz to 100 kHz, with potential amplitude of 10 mV. The direct potential during measurements was set to 0 and all the measurements were done versus an open circuit potential. The open-source modeling program, EIS Spectrum Analyzer (Bondarenko and Ragoisha 2005) was used to model the impedance spectra with an equivalent Randles circuit. The program is available at http://www.abc.chemistry.bsu.by/vi/analyser/.

### 2.6. Scanning electron microscopy

Electrodes were mounted on aluminum stubs (32mm diameter) with carbon adhesive discs (Agar Scientific, Oxford Instruments SAS, Gomez-la-Ville, France) and visualized by field emission gun scanning electron microscopy (SEM FEG) as secondary and backscattered electrons images (2 keV, spot size 30) under high vacuum conditions with a Hitachi SU5000 instrument (Milexia, Saint-Aubin, France). Sample preparation and Scanning Electron Microscopy analyses were performed at the Microscopy and Imaging Platform MIMA2 (INRAE, Jouy-en-Josas, France) DOI: MIMA2, INRAE, 2018. Microscopy and Imaging Facility for Microbes, Animals and Foods, https://doi.org/10.15454/1.5572348210007727E12.

## 3. Results and Discussion

### 3.1. Gold-leaf electrode fabrication

We prepared the GLE on a low cost PVC foil by assembling four PVC sheets to obtain an easy to handle solid support (Fig. 1a). To make this support hydrophobic, an insulating layer was created by spraying PTFE on the upper sheet. This step was performed in order to prevent wettability of aqueous samples during measurements. Finally, two gold leaves were deposited on the top. Both, one or two gold leaves can be used to fabricate GLE, but we found that two leaves gave a more homogeneous surface. After assembling all layers by lamination, the GLE platform was fabricated using laser ablation creating a 2.5 mm diameter working electrode zone. The dimensions of fabricated working and counter electrodes are presented in Fig. 1b. This design enables an electrochemical activity using a small sample volume (50 μL). Fig. 1c and 1d show the assembled layers before and after laser ablation, respectively, indicating that at least 18 electrodes can be fabricated through a single production process. 2D and 3D profiles of the working electrode are presented in Figs. 1e and 1f to illustrate the gold surface roughness. The surface of the GLE showed regular undulating features with micrometer-scale hills and valleys.

We estimated that our home-made GLE is order of magnitude less expensive than commercial gold SPE taking into account that affordable pure gold leaf sheets cost €50 / 25, and that at least 18 electrodes can be obtained from a single gold sheet. Therefore the fabrication method seems suitable for use in both standard labs and low-resource settings.

### 3.2. Gold-leaf electrode characterization

Cyclic voltammetry was performed in sulfuric acid to characterize the crystallinity of GLE. The CV shows a single oxidation peak at +1.3 V (vs. Ag/AgCl) along with a small oxidation shoulder, with the reduction peak at +0.80 V (vs. Ag/AgCl), corresponding to the electrochemical reduction of the gold oxide layer formed in the forward scan (Fig. 2a). Such a CV curve is typical for crystalline bare gold with a homogeneous surface (Sukeri et al. 2015).

**Figure 2.**
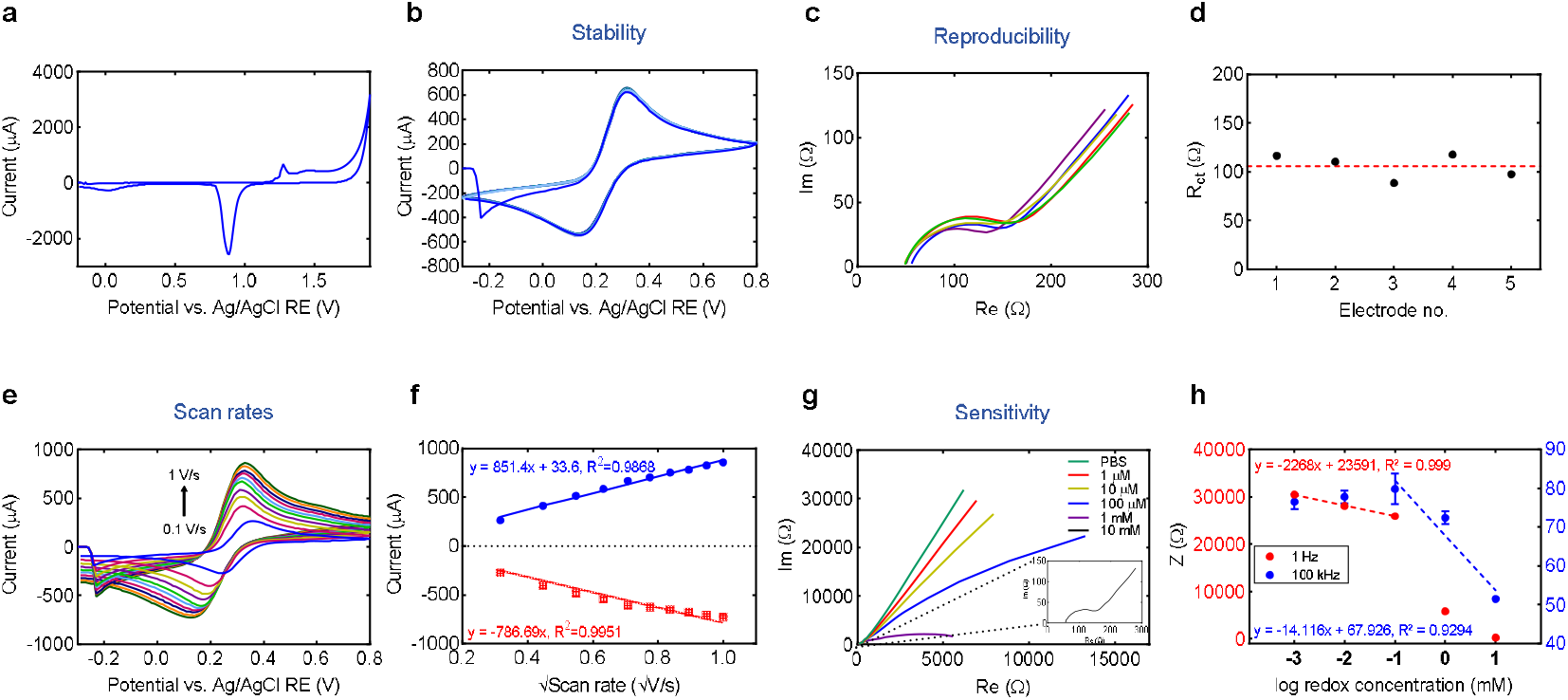
GLE characterization. (a) CV of 0.5 M H_2_SO_4_ at the scan rate of 0.5 V/s. (b) CV of 1 mM K_3_[Fe(CN)_6_]/K_4_[Fe(CN)_6_] in PBS recorded 25 successive times using the same GLE. (c) EIS responses obtained with five different GLEs for 10 mM K_3_[Fe(CN)_6_]/K_4_[Fe(CN)_6_] in PBS. (d) Reproducibility of the charge transfer resistance, R_ct_, calculated after fitting curves in (c) for different electrodes. (e) CV of 10 mM K_3_[Fe(CN)_6_]/K_4_[Fe(CN)_6_] in PBS at scan rates from 0.1 to 1 V/s recorded using GLE. (f) The dependence of 10 mM K_3_[Fe(CN)_6_]/K_4_[Fe(CN)_6_] anionic and cationic peak intensity on the square root of scan rate. (g) EIS responses obtained with different concentrations of the K_3_[Fe(CN)_6_]/K_4_[Fe(CN)_6_] redox probe in PBS. (h) Impedance module variation at 1 Hz and 100 kHz with the respect to K_3_[Fe(CN)_6_]/K_4_[Fe(CN)_6_] concentration with linear fitting of values in range 0.001 mM to 0.1 mM and 0.1 mM to 10 mM, respectively.

The electrode was next characterized for its stability and reproducibility which are always a concern with a house-made material and device. Multiple CV scans were performed using a 10 mM redox probe K_3_[Fe(CN)_6_]/K_4_[Fe(CN)_6_] in PBS. The obtained voltammograms in Fig. 2b show that the Fe^2+^/Fe^3+^ redox reaction displayed a well-defined oxidation and reduction peaks. Their intensities had constant values after 25 successive scans indicating a very good stability of the GLE. Using the Randles–Sevcik equation (Eq. 1), the effective surface area of GLE was estimated to be 14.2 mm^2^, with the R_f_ of 0.72 (Fig. S2). For comparison, the commercial DropSens 220AT electrode, has the effective area of about 8.4 mm^2^ and the R_f_ of 0.66 (Fig. S2). It is important to mention that the mass transfer is a limiting factor in an electrochemical measurement of effective area, since the inner part of the electrode is not effected by the electroactive compound due to the its reaction in the upper part of the electrode (Krejci et al. 2014). However, as expected, the GLE with higher surface R_*f*_ gave higher sensitivity for the Fe^2+^/Fe^3+^ redox reaction than the commercial electrode (Fig. S2).

EIS was used to assess the reproducibility of electrodes in a solution of the Fe^2+^/Fe^3+^ redox couple in PBS (Fig. 2c). The impedance spectra consisted of a semi-circle portion with the diameter calculated as electron transfer resistance and a linear portion which represents a diffusion controlled process. Excellent reproducibility was obtained using five different electrodes as all analyzed devices displayed a very similar response with a low variability especially in the semi-circle portion (Figs. 2c and 2d).

Next, the GLEs were characterized with CV using the ferri/ferrocyanide redox probe in PBS at varying speeds of polarization. As shown in Fig. 2e, the current increased with increasing scan rate. The intensity of both the anion oxidation peak and cation reduction peak of the redox probe plotted against the square root of the scan rate indicated linear relationships (Fig. 2f). The linearity suggests that diffusion controlled the oxidation/reduction of K_3_[Fe(CN)_6_]/K_4_[Fe(CN)_6_] at the GLE while stability of the faradaic peak potentials suggests rapid kinetics of the probe oxidation/reduction.

Finally, the dependence of the ferri/ferrocyanate impedance signal on its concentration in PBS buffer was evaluated (Fig. 2g). A linear range of the impedance module was observed at 1Hz and 100 kHz for concentrations ranging from 0.001 mM to 0.1 mM and 0.1 mM to 10 mM, respectively (Fig. 2h). This shows that the electrode enables a reliable and sensitive impedance detection of ferri/ferrocyanate with settings from 1 Hz to 100 kHz.

### 3.3. Immuno-sensor construction

To demonstrate the capability of GLE for applications in biosensor development, we examined immuno-detection of *E. coli* cells. A sensitive, low-cost and rapid detection platform for *E. coli* is expected by many professionals in health and water/food safety industries because the bacterium, when ingested, even in low doses can cause vomiting, bloody diarrhea and severe abdominal cramps.

The GLE was functionalized with a specific capturing antibody which recognizes the whole cell of *E. coli* in order to enable bacterial detection without preliminary cell lysis (Fig. 3a). To attach the antibody to the electrode, the surface was modified with MPA that formed a self-assembled monolayer on the gold. After thorough cleansing and immersion into MPA ethanolic solution, the GLE was characterized by water contact angles. The pixel images in Fig. 3b show that the initial contact angle values of a PBS droplet decreases from 70.7(4)° to 53.53(27)° after MPA assembling on the surface. These values indicate the increase in surface hydrophobicity after modification and are in agreement with already published results (Braiek et al. 2012; He et al. 2009).

**Figure 3.**
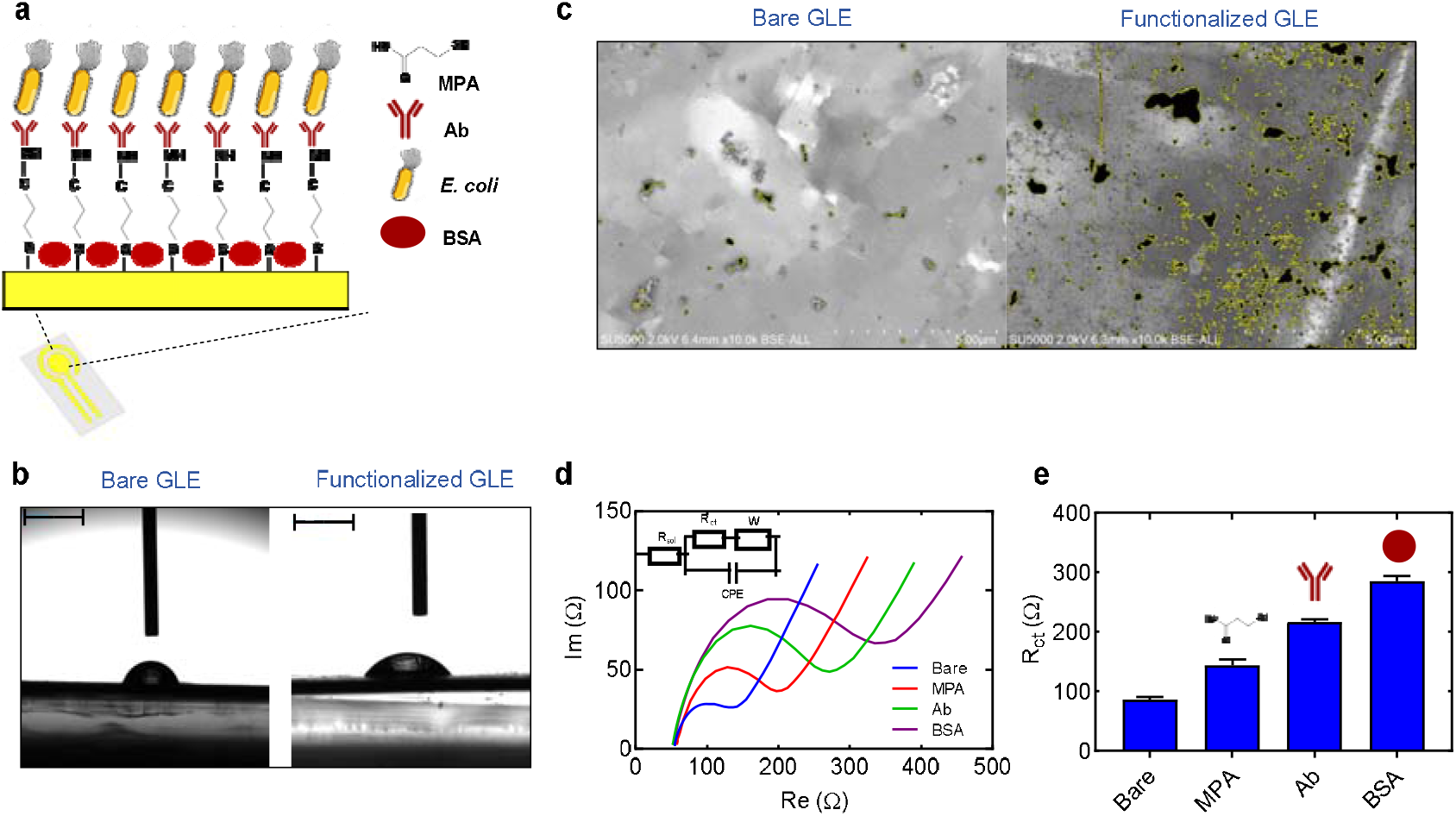
Electrode functionalization. (a) Schematic illustration of the immuno-sensor construction for *E. coli* detection. (b) Contact angles and properties of MPA, where (left image) bare gold with the angle measured at 70.7 °, and (right image) MPA monolayer with the angle measured at 53.53°. Scale bar, 2 mm. (c) SEM images of the bare GLE, and GLE functionalized with MPA and anti-*E. coli* antibody. Note that the sizes of visualized areas corresponding to organic compounds attached to the electrode surface increased significantly after functionalization. (d) Nyquist plot of impedance spectra obtained on GLE bare and GLE modified with MPA, antibody and BSA in 10 mM K_3_[Fe(CN)_6_]/K_4_[Fe(CN)_6_] in PBS at a frequency range between 1Hz and 100 kHz. Inset, equivalent electrical circuit applied to fit the Nyquist plot. (e) The fitting values of the charge transfer resistance, R_ct_, at various steps of GLE functionalization.

The antibody was immobilized on the functionalized electrode via an amino link after activation of the carboxylic acid-terminated MPA monolayer with EDC/NHS solution (Fig. 3a). Finally, the surface was blocked with BSA to prevent nonspecific binding. To assess the quantitative analysis of the functionalization efficiency, we performed SEM visualization of the electrode surface before and after antibody immobilization and blocking (Fig. 3c). The SEM image of a bare GLE showed a surface of a lighter color compared to that of the functionalized electrode indicating that low electro-charged biological molecules were immobilized (Fig. S3). The SEM images for modified electrodes showed a random distribution of dark spots when observed in back-scattered secondary electron mode that corresponded to organic molecules bound to the gold surface (Fig. 3c and Fig. S3). To further validate GLE surface modification, the ferri/ferrocyanide redox probe in PBS was monitored after each functionalization step. EIS monitoring of electrode functionalization showed a significant increase in semicircle diameter in the Nyquist diagrams after different stages of immuno-sensor construction confirming formation of different layers on the electrode surface (Fig. 3d and e).

### 3.4. Application of gold-leaf microelectrode for E. coli detection

The GLE carrying the anti-*E. coli* antibody was then exposed to increasing concentrations of *E. coli* cells in PBS. The corresponding Nyquist plots of the impedance spectra are reported in Fig. 4a. The impedance measurements were fitted using an equivalent circuit of the Randles cell shown in the inset of Fig. 4b and the corresponding electronic circuit elements were calculated. In brief, the circuit includes four elements: the ohmic resistance of electrolyte solution, R_sol_; the constant phase element, CPE, reflecting the non-homogeneity of the layer in the presence of large entities; the Warburg impedance, Z_w_, which is related to the low frequency part of the spectrum, where the diffusion mechanism controls the measured impedance and charge transfer resistance R_ct_. Of these elements, R_ct_ and CPE depend on the dielectric and insulating features at the electrode/electrolyte interface, and reflect modification of the electrode surface due to target capturing by the antibody on the electrode surface. The major change was an increase of the charge transfer resistance R_ct_ that is related to the mass transfer phenomenon and/or the dielectric or conductive properties of the captured bacterial cells. The charge transfer resistance was, thus, selected to quantify bacterial cell detection. R_ct_ showed increasing trends with an increasing amount of *E. coli* cells and a good linear dependence in the range of 10^1^ to 10^7^ CFU/ml (Fig. 4b). The regression equation was *y* = 78.837 (*x*) + 367.75, with R^2^ of 0.9862. The experimental limit of detection (LoD) was 10 CFU/ml while the calculated LoD was 2 CFU/ml (according to IUPAC rules with the formula (3*S/d*) where *S* is the standard deviation of a blank test measured with 3 biosensors and *d* is the sensitivity deduced from the slope of the linear curve). Such low LoD is comparable with values obtained for *E. coli* detection using advanced electrochemical biosensors as listed in Table 1, recently proposed in the literature. Moreover, detection using GLE is by five orders of magnitude more sensitive than the traditional immunological ELISA test (Galikowska et al. 2011). The obtained low LoD corroborates the clinically relevant infectious doses of *E. coli* (100 cells), which suggests that the immuno-sensor based on GLE can be used without further improvement of its sensitivity.

**Figure 4.**
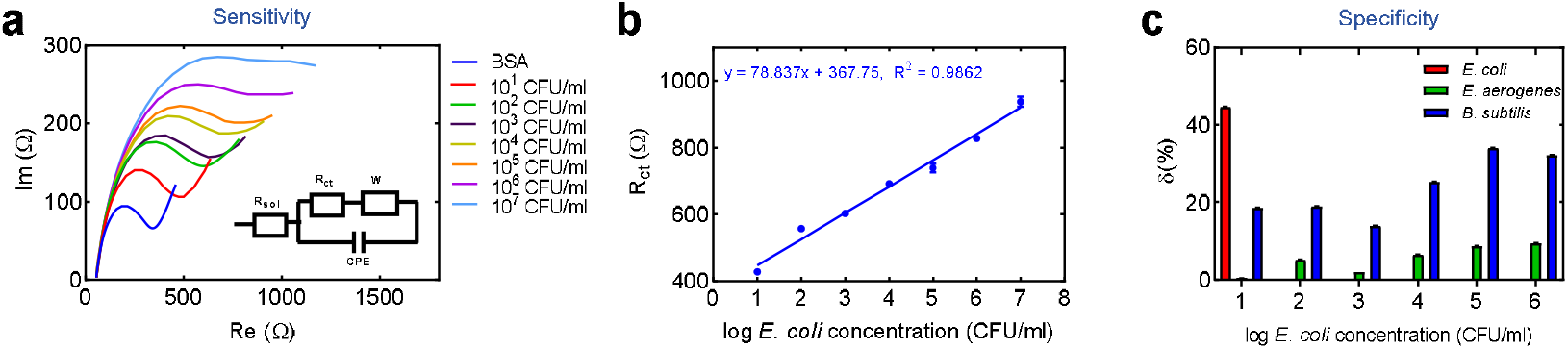
Electrochemical detection of *E. coli*. (a) Nyquist plot of impedance spectra obtained from the increasing amount of *E. coli* cells from 10^1^ to 10^7^ CFU/ml in PBS and 10 mM K_3_[Fe(CN)_6_]/K_4_[Fe(CN)_6_]. Inset, equivalent electrical circuit applied to fit the Nyquist plot. (b) Calibration plots showing the change of charge transfer resistance, R_ct_, as a function of the different concentrations of *E. coli*. Data points are the mean values obtained in 3 independent experiments ± SD. (c) Positive-to-negative response ratio of the GLE-based immune-sensor for *E. coli*, *E. aerogenes* and *B. subtilis* (δ = [R_ct(sample)_ - R_ct(immobilized antibody)_ / R_ct(immobilized antibody)_] × 100%).

**Table 1.** Performance comparison of the designed GLE-biosensor with previously reported electrochemical biosensors for *E. coli*, reported in literature.

| Detection method | Dynamic range (CFU/ml) | LoD (CFU/ml) | Reference |
| --- | --- | --- | --- |
| Square wave voltammetry | $10^1-10^4$ | 10 | (Bonnet et al. 2018) |
| Impedance | $3 \times 10^2-3 \times 10^5$ | 300 | (Dastider et al. 2013) |
| Micro-conductometry | $10^1-10^5$ | 10 | (El Ichi et al. 2014) |
| Amperometry | $10^2-10^7$ | 100 | (Lin et al. 2008) |
| Amperometry | $10^2-10^8$ | 100 | (Laczka et al. 2011) |
| Voltammetry | $4.8 \times 10^1-10^5$ | 48 | (Safavieh et al. 2012) |
| Voltammetry | $4.8 \times 10^5-10^7$ | $4.8 \times 10^5$ | (Varshney and Li 2007) |
| Impedance | $10^1-10^4$ | 10 | (Maalouf et al. 2007) |
| Impedance | $10-10^4$ | 2 | (Dos Santos et al. 2013) |
| Square-wave voltammetry | $10^1-10^7$ | 10 | (Gunasekaran et al. 2022) |
| Voltammetry | $10^1-10^7$ | 10 | (Khan et al. 2021) |
| Impedance | $10^1-10^7$ | 2 | Our work |

The sensor selectivity was studied on *E. aerogenes* (Gram-negative) and *B. subtilis* (Gram-positive) as nonspecific bacterial targets for the capturing antibody. The control bacterial cells were introduced in the GLE sensing area using the same experimental protocol used for *E. coli* (Fig. 4c). The relative change in the R_ct_ caused by various bacterial concentrations versus R_ct_ of BSA, δ, indicated that no significant response was detected for both control bacteria in the concentration range from 10^1^ CFU/ml to 10^6^ CFU/ml compared to the signal increase observed with 10^1^ CFU/ml of *E. coli*. Thus, the method demonstrates acceptable sensor specificity.

## 4. Conclusions

In this work, we have presented a new design and fabrication method of a gold leaf electrode and a highly performant electrochemical immunosensor for detection of *E. coli* cells. The key issues in developing disposable electrodes that can be easily used for rapid, sensitive and low cost biosensing applications remain the demand for low sample volume, high conductivity and large surface area of the working electrode. Exploiting 24-karat gold leaf as a material for electrode fabrication, the sensing platform was designed using hot lamination and laser ablation. The manufacturing process is affordable as it does not require any support layers or masks nor a cleanroom or a postprocessing. In addition, it is very fast: in laboratory conditions one electrode is fabricated in less than a minute, while technologies such as lithography, screen, or inkjet printing require a couple of hours. Moreover, multiple copies of the electrode can be manufactured in a single run, with the production cost of only about €0.1 per copy. Due to the narrow laser beam and its resolution, high repeatability and reproducibility can be achieved. Moreover, a complex electrode geometry can be realized with a very small resolution and narrow spacing between electrodes.

Innovations in biosensing have not only upgraded healthcare, agriculture, food and environmental safety measures to reduce hazards but have also economically benefited industries. The potential of GLEs was demonstrated in the realization of an immunosensor for the detection of *E. coli*. With the detection limit of 2 CFU/ml and a linear dynamic range of 10 – 10^7^ CFU/ml the sensor based on GLE outperforms most reported devices that use a more complex surface chemistry and functionalization to enhance the signal and/or require multifaceted fabrication. Taking into account that *E. coli* is of great importance to the quality control of food and water and that traditional methods are not suitable for direct and rapid bacterial detection (Cossettini et al. 2022; Kotsiri et al. 2022) the presented sensitive biosensing of *E. coli* may play a significant role in the field of on-site analysis and monitoring.

## CRediT authorship contribution statement

**Ivana Podunavac:** Conceptualization, Methodology, Formal analysis, Investigation. **Manil Kukkar:** Conceptualization, Methodology. **Vincent Léguillier:** Investigation, Writing - original draft. **Francesco Rizzotto:** Formal analysis, Investigation. **Zoran Pavlovic:** Methodology, Investigation, Visualization. **Ljiljana Janjušević:** Methodology, Inversigation. **Vlad Costache:** Investigation, Formal analysis. **Vasa Radonic:** Supervision, Conceptualization, Methodology, Writing - review & editing. **Jasmina Vidic:** Supervision, Conceptualization, Methodology, Writing - original draft, Writing - review & editing.

## Acknowledgments

This work was supported in part by the IPANEMA project, which received funding from the European Union’s Horizon 2020 research and innovation programme under grant agreement N° 872662, and in part by the French National Agency for Research (ANR-21-CE21-0009 SIENA project, to JV), OneHealth2021 (to JV), Science Fund of the Republic of Serbia (#Grant MircoLabAptaSense, N° 7750276, to VR). We thank Maria Vesna Nikolic (IMSI, Serbia) for discussion and critical reading of the manuscript and the MIMA2 platform for access to electron microscopy equipment (MIMA2, INRAE, 2018. Microscopy and Imaging Facility for Microbes, Animals and Foods, https://doi.org/10.15454/1.5572348210007727E12MIMA2).

## References

Bernalte, E., Marín-Sánchez, C., Pinilla-Gil, E., Brett, C.M., 2013. Characterisation of screen-printed gold and gold nanoparticle-modified carbon sensors by electrochemical impedance spectroscopy. Journal of Electroanalytical Chemistry 709, 70–76.

Bobrinetskiy, I., Radovic, M., Rizzotto, F., Vizzini, P., Jaric, S., Pavlovic, Z., Radonic, V., Nikolic, M.V., Vidic, J., 2021. Advances in nanomaterials-based electrochemical biosensors for foodborne pathogen detection. Nanomaterials 11(10), 2700.

Bondarenko, A., Ragoisha, G., 2005. Progress in chemometrics research. Nova Science, New York,(http://www.abc.chemistry.bsu.by/vi/), 1110–1112.

Bonnet, R., Farre, C., Valera, L., Vossier, L., Léon, F., Dagland, T., Pouzet, A., Jaffrézic-Renault, N., Fareh, J., Fournier-Wirth, C., 2018. Highly labeled methylene blue-ds DNA silica nanoparticles for signal enhancement of immunoassays: application to the sensitive detection of bacteria in human platelet concentrates. Analyst 143(10), 2293–2303.

Braiek, M., Rokbani, K.B., Chrouda, A., Mrabet, B., Bakhrouf, A., Maaref, A., Jaffrezic-Renault, N., 2012. An electrochemical immunosensor for detection of Staphylococcus aureus bacteria based on immobilization of antibodies on self-assembled monolayers-functionalized gold electrode. Biosensors 2(4), 417–426.

Cossettini, A., Vidic, J., Maifreni, M., Marino, M., Pinamonti, D., Manzano, M., 2022. Rapid detection of Listeria monocytogenes, Salmonella, Campylobacter spp., and Escherichia coli in food using biosensors. Food Control, 108962.

Dastider, S.G., Barizuddin, S., Dweik, M., Almasri, M., 2013. A micromachined impedance biosensor for accurate and rapid detection of E. coli O157: H7. RSC advances 3(48), 26297–26306.

Dos Santos, M.B., Agusil, J., Prieto-Simón, B., Sporer, C., Teixeira, V., Samitier, J., 2013. Highly sensitive detection of pathogen Escherichia coli O157: H7 by electrochemical impedance spectroscopy. Biosensors and Bioelectronics 45, 174–180.

El Ichi, S., Leon, F., Vossier, L., Marchandin, H., Errachid, A., Coste, J., Jaffrezic-Renault, N., Fournier-Wirth, C., 2014. Microconductometric immunosensor for label-free and sensitive detection of Gramnegative bacteria. Biosensors and bioelectronics 54, 378–384.

Galikowska, E., Kunikowska, D., Tokarska-Pietrzak, E., Dziadziuszko, H., Łoś, J.M., Golec, P., Węgrzyn, G., Łoś, M., 2011. Specific detection of Salmonella enterica and Escherichia coli strains by using ELISA with bacteriophages as recognition agents. European journal of clinical microbiology & infectious diseases 30(9), 1067–1073.

Gunasekaran, D., Gerchman, Y., Vernick, S., 2022. Electrochemical Detection of Waterborne Bacteria Using Bi-Functional Magnetic Nanoparticle Conjugates. Biosensors 12(1), 36.

He, F., Shen, Q., Jiang, H., Zhou, J., Cheng, J., Guo, D., Li, Q., Wang, X., Fu, D., Chen, B., 2009. Rapid identification and high sensitive detection of cancer cells on the gold nanoparticle interface by combined contact angle and electrochemical measurements. Talanta 77(3), 1009–1014.

Khan, S., Akrema, Qazi, S., Ahmad, R., Raza, K., Rahisuddin, 2021. In Silico and electrochemical studies for a ZnO–CuO-based immunosensor for sensitive and selective detection of E. coli. ACS omega 6(24), 16076–16085.

Kotsiri, Z., Vantarakis, A., Rizzotto, F., Kavanaugh, D., Ramarao, N., Vidic, J., 2019. Sensitive Detection of E. coli in Artificial Seawater by Aptamer-Coated Magnetic Beads and Direct PCR. Applied Sciences 9(24), 5392.

Kotsiri, Z., Vidic, J., Vantarakis, A., 2022. Applications of biosensors for bacteria and virus detection in food and water–A systematic review. journal of environmental sciences 111, 367–379.

Krejci, J., Sajdlova, Z., Nedela, V., Flodrova, E., Sejnohova, R., Vranova, H., Plicka, R., 2014. Effective surface area of electrochemical sensors. Journal of the Electrochemical Society 161(6), B147.

Laczka, O., Maesa, J.-M., Godino, N., del Campo, J., Fougt-Hansen, M., Kutter, J.P., Snakenborg, D., Muñoz-Pascual, F.-X., Baldrich, E., 2011. Improved bacteria detection by coupling magneto-immunocapture and amperometry at flow-channel microband electrodes. Biosensors and Bioelectronics 26(8), 3633–3640.

Lin, Y.-H., Chen, S.-H., Chuang, Y.-C., Lu, Y.-C., Shen, T.Y., Chang, C.A., Lin, C.-S., 2008. Disposable amperometric immunosensing strips fabricated by Au nanoparticles-modified screen-printed carbon electrodes for the detection of foodborne pathogen Escherichia coli O157: H7. Biosensors and Bioelectronics 23(12), 1832–1837.

Maalouf, R., Fournier-Wirth, C., Coste, J., Chebib, H., Saïkali, Y., Vittori, O., Errachid, A., Cloarec, J.-P., Martelet, C., Jaffrezic-Renault, N., 2007. Label-free detection of bacteria by electrochemical impedance spectroscopy: comparison to surface plasmon resonance. Analytical chemistry 79(13), 4879–4886.

Manges, A.R., Geum, H.M., Guo, A., Edens, T.J., Fibke, C.D., Pitout, J.D., 2019. Global extraintestinal pathogenic Escherichia coli (ExPEC) lineages. Clinical microbiology reviews 32(3), e00135–00118.

Marin, M., Nikolic, M.V., Vidic, J., 2021. Rapid point-of-need detection of bacteria and their toxins in food using gold nanoparticles. Comprehensive Reviews in Food Science and Food Safety 20(6), 5880–5900.

Matsui, Y., Hamamoto, K., Kitazumi, Y., Shirai, O., Kano, K., 2017. Diffusion-controlled mediated electron transfer-type bioelectrocatalysis using microband electrodes as ultimate amperometric glucose sensors. Analytical Sciences 33(7), 845–851.

Prasertying, P., Jantawong, N., Sonsa-Ard, T., Wongpakdee, T., Khoonrueng, N., Buking, S., Nacapricha, D., 2021. Gold leaf electrochemical sensors: applications and nanostructure modification. Analyst 146(5), 1579–1589.

Safavieh, M., Ahmed, M.U., Tolba, M., Zourob, M., 2012. Microfluidic electrochemical assay for rapid detection and quantification of Escherichia coli. Biosensors and Bioelectronics 31(1), 523–528.

Singh, M., Haverinen, H.M., Dhagat, P., Jabbour, G.E., 2010. Inkjet printing—process and its applications. Advanced materials 22(6), 673–685.

Sui, Y., Zorman, C.A., 2020. Inkjet printing of metal structures for electrochemical sensor applications. Journal of The Electrochemical Society 167(3), 037571.

Sukeri, A., Saravia, L.P.H., Bertotti, M., 2015. A facile electrochemical approach to fabricate a nanoporous gold film electrode and its electrocatalytic activity towards dissolved oxygen reduction. Physical Chemistry Chemical Physics 17(43), 28510–28514.

Tang, Y., Li, Z., Luo, Q., Liu, J., Wu, J., 2016. Bacteria detection based on its blockage effect on silicon nanopore array. Biosensors and Bioelectronics 79, 715–720.

Varshney, M., Li, Y., 2007. Interdigitated array microelectrode based impedance biosensor coupled with magnetic nanoparticle–antibody conjugates for detection of Escherichia coli O157: H7 in food samples. Biosensors and Bioelectronics 22(11), 2408–2414.

Vidic, J., Manzano, M., 2021. Electrochemical biosensors for rapid pathogen detection. Current Opinion in Electrochemistry 29, 100750.

Vidic, J., Pla-Roca, M., Grosclaude, J., Persuy, M.-A., Monnerie, R., Caballero, D., Errachid, A., Hou, Y., Jaffrezic-Renault, N., Salesse, R., 2007. Gold surface functionalization and patterning for specific immobilization of olfactory receptors carried by nanosomes. Analytical Chemistry 79(9), 3280–3290.

Zamani, M., Yang, V., Maziashvili, L., Fan, G., Klapperich, C.M., Furst, A.L., 2021. Surface requirements for optimal biosensing with disposable gold electrodes. ACS measurement science Au 2(2), 91–95.

Zhang, J., Jiang, Y., Xia, X., Wu, J., Almeida, R., Eda, S., Qi, H., 2020. An on-site, highly specific immunosensor for Escherichia coli detection in field milk samples from mastitis-affected dairy cattle. Biosensors and Bioelectronics 165, 112366.

