## Supplementary data for "Low-cost gold-leaf electrode as a platform for *Escherichia coli* immuno-detection"

84its geometric area was 19.62 mm^2^

**Table of Contents:**

**Figure S1………………………………………………………………………………….2**

**Figure S2………………………………………………………………………….……....3**

**Figure S3………………………………………………………………………………….4**

### 1. Experimental setup

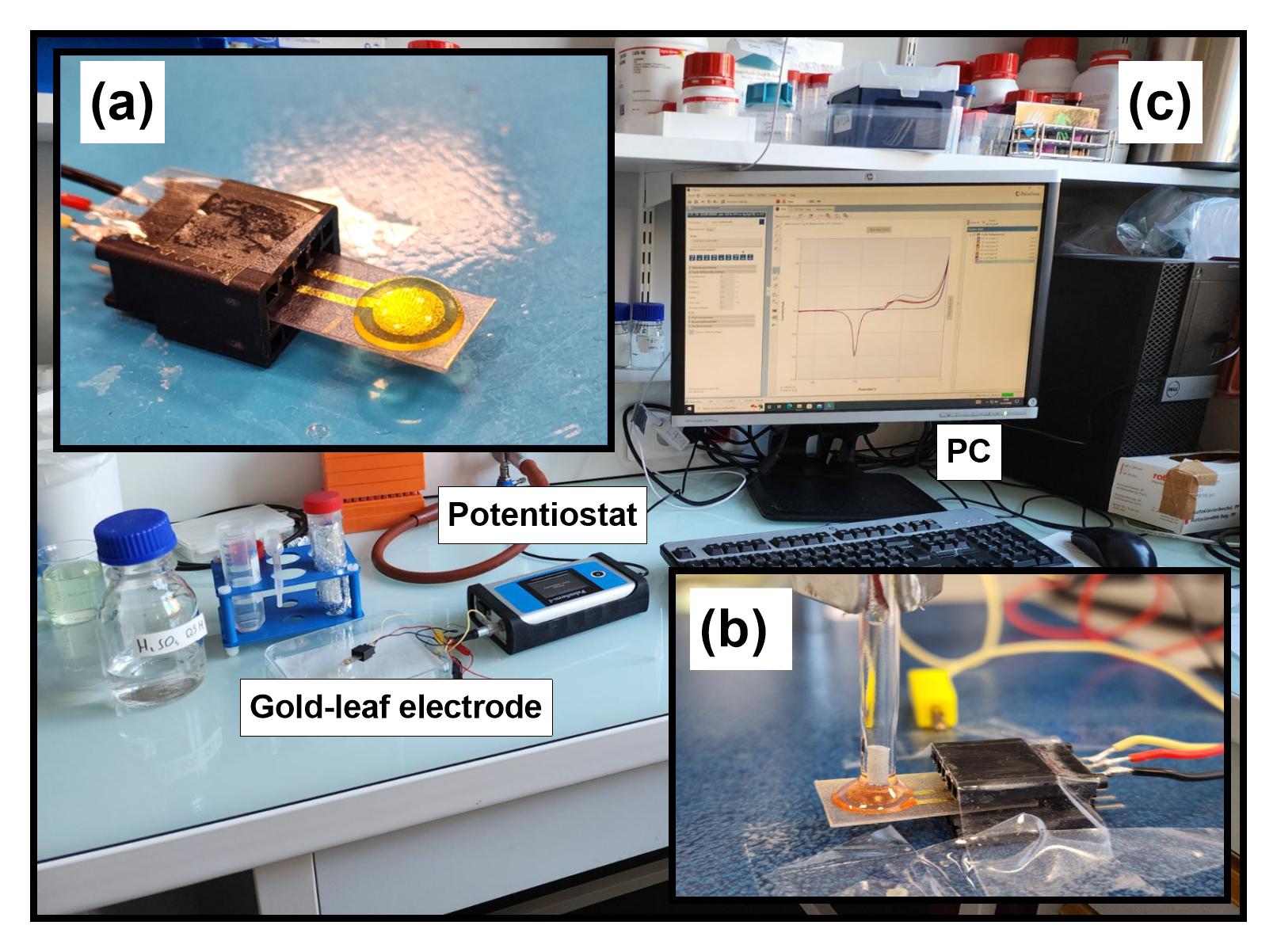

Figure S1. Photo images of the gold leaf electrode (GLE) and GLE-based sensor. (a) GLE used in a two-electrode biosensor, (b) GLE associated with a commercial Ag/AgCl referent electrode used in a three-electrode biosensor, (c) Experimental setup.

### 2. Comparison of roughness of in-house gold leaf and commercial Dropsens 220AT electrodes

In order to determinate the effective surface area of the electrode via the Randles–Sevcik equation, CV measurements for 10 mM K_4_[Fe(CN)_6_]/ K_3_[Fe(CN)_6_] in PBS with scan rate ranging from 0.1 V/s to 1 V/s were performed for both commercial DropSens 220AT electrode and our in-house GLE.

From Eq. 1 and slopes in Fig. S2c and Fig. S2d, the electrode effective area was calculated to be 14.2 mm^2^ and 8.4 mm^2^ for GLE and Dropsens 220AT electrode, respectively. By comparing their effective and geometric area (also known as roughness factor), the roughness factor of the GLE electrode is about 91 % higher than that of the Dropsens 220AT electrode (R_fGLE_/R_f220AT_ x 100). The high R_f_ of GLE is in agreement with the observed 3D profile of the GLE surface in Fig. 1f showing significant micrometer-scale electrode roughness.

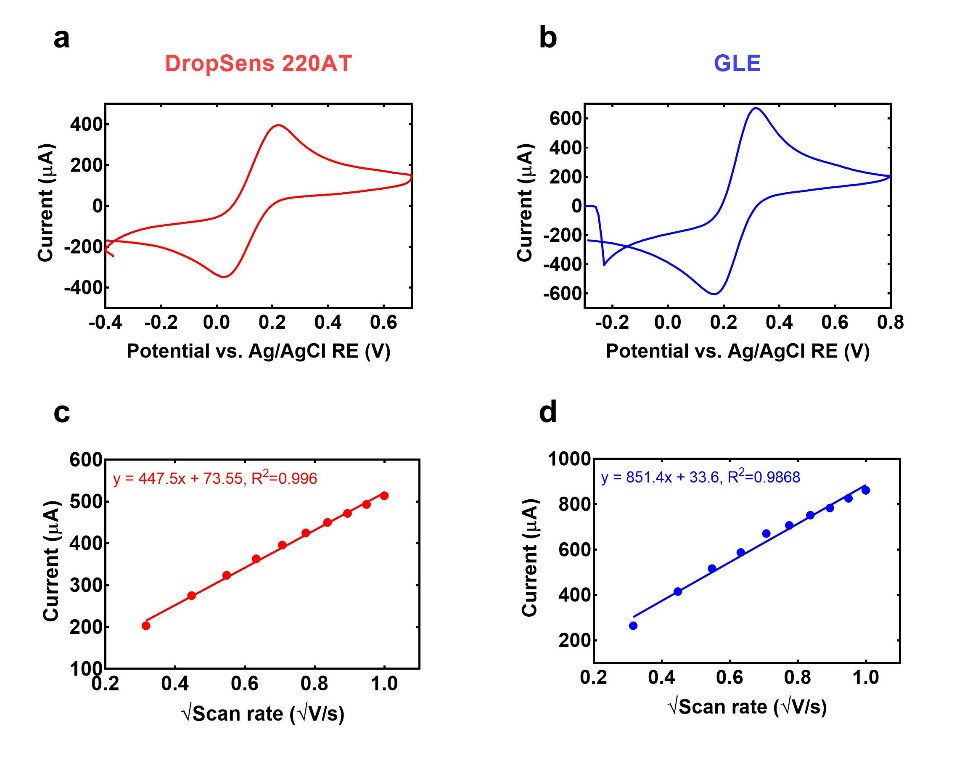

Figure S2. (a) CV of 10 mM K_4_[Fe(CN)_6_]/ K_3_[Fe(CN)_6_] in PBS at DropSens 220AT electrode at scan rate 0.5 V/s; (b) CV of 10 mM K_4_[Fe(CN)_6_]/ K_3_[Fe(CN)_6_] in PBS at GLE electrode at scan rate 0.5 V/s; (c) The dependance of 10 mM K_4_[Fe(CN)_6_]/ K_3_[Fe(CN)_6_] oxidation peak intensity on the square root of scan rate for DropSens 220AT electrode. (d) The dependance of 10 mM K_4_[Fe(CN)_6_]/ K_3_[Fe(CN)_6_] oxidation peak intensity on the square root of scan rate for GLE electrode.

### 3. SEM of bare and functionalized gold leaf electrode

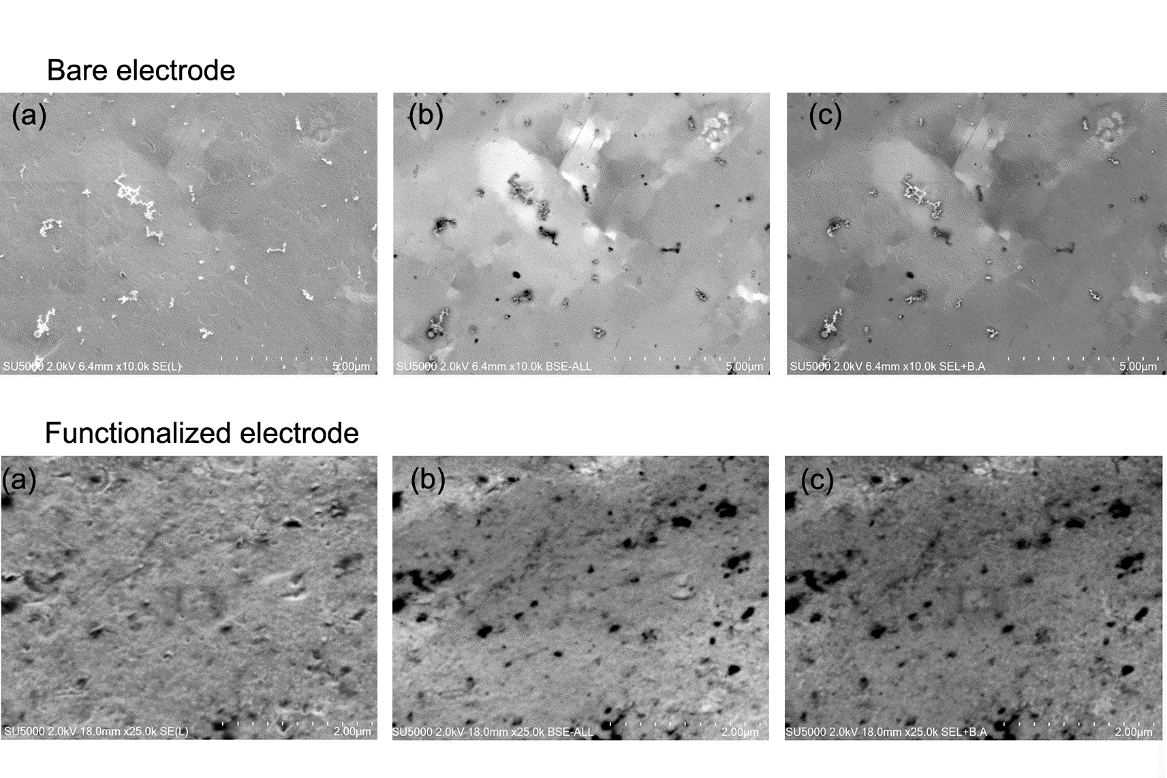

Figure S3. SEM images of the working electrodes of GLEs before (bare electrode) and after immobilization of MPA (functionalized electrode). (a) Secondary electron (SE) mode to observe the topography of the surface, (b) Backscattered secondary electron (BSE) mode to reveal the internal structure of the surface, and (c) merged SE and BSE.
